# Two diseases, same person: moving towards a combined HIV and TB continuum of care

**DOI:** 10.1101/186833

**Authors:** Reuben Granich, Somya Gupta

**Author notes:** Address correspondence and reprint requests to: Reuben Granich, MD, MPH, 4448 Springdale St., NW, Washington DC 20016.

## Abstract

**Setting:** The Human Immunodeficiency Virus (HIV) and *Mycobacterium tuberculosis* syndemic remains a global public health threat. Separate HIV and TB global targets have been set, however, success will depend on achieving combined disease control objectives and care continua.

**Objective:** Review available policy, budgets and data to re-conceptualize TB and HIV disease control objectives by combining HIV and TB care continua.

**Methods:** For 22 WHO TB and TB/HIV priority countries, we used 2014 and 2015 data from the HIV90-90-90watch website, UNAIDS Aidsinfo, and WHO 2016 Global TB Report. Global resources available in TB and HIV/TB activities for 2003-2017 was collected from publically available sources.

**Results:** In 22 high burden countries people living with HIV (PLHIV) on ART ranged from 9-70%; viral suppression was 38-63%. TB treatment success ranged from 34-94% with 13 (43% HIV/TB burden) countries above 80% TB treatment success. From 2003-2017, global international and domestic resources for HIV-associated TB and TB averaged $2.6 billion per year; the total for 2003-2017 was 39 billion dollars.

**Conclusion:** Reviewing **c**ombined HIV and TB targets demonstrate disease control progress and challenges. Using an integrated HIV and TB continuum supports HIV and TB disease control efforts focused on improving both individual and public health.

**Funding:** None

## Introduction

The Human Immunodeficiency Virus (HIV) and tuberculosis (TB) syndemic continues to be a major global public health threat. In 2015, 2 billion people were living with *Mycobacterium tuberculosis* infection and 10.4 million developed TB disease. Of these, 1.2 million (11%) were people living with HIV (PLHIV) (1). Of the 580,000 multi-drug resistant (MDR) tuberculosis cases, 92,000 (16%) received treatment and only 52% reported treatment success--extensively drug-resistant (XDR) TB fared worse, reporting a 28% treatment success rate in 2013 (1). Many of these deadly, airborne, TB cases occur among highly vulnerable PLHIV who, without HIV treatment, have a 10% risk of developing TB per year (2). Antiretroviral treatment (ART) reduces the individual risk of TB by approximately 65% (3), and by 90% when coupled with isoniazid preventive therapy (IPT) (4, 5). However, by mid-2016, only 18.2 million (50%) of PLHIV were on HIV treatment and most were not offered IPT (6). Additionally, HIV-associated TB is curable with early treatment for TB and HIV (7, 8). However, 2015 global TB case detection was only 59% (1). Although there has been impressive improvements in both HIV and TB treatment coverage, the gap contributed to millions of new TB infections, 1.4 million TB deaths (in HIV-negative), 2.1 million HIV infections, 1.1 million HIV-associated deaths, and 1.2 million HIV-associated TB cases and 390,000 HIV-associated TB deaths (1, 6).

Each new HIV-associated TB case among people who are not on ART is a missed public health opportunity or failure. To prevent HIV and TB illness, death and transmission, the Joint United Nations Program on HIV/AIDS (UNAIDS) proposed the 90-90-90 target by 2020 to end AIDS (9)--diagnose 90% of people living with HIV, provide ART for 90% of people diagnosed, and 90% of people on ART with viral suppression. The target will be attained when at least 29.5 (73%) million PLHIV are on ART and virally suppressed (17 million in sub-Saharan Africa alone) (9). The 90-90-90 target envisions 27% of PLHIV unsuppressed which is further reduced to 14% unsuppressed for the 2030 target of 95-95-95 (9). For ending TB, the Stop TB Partnership has issued a Global Plan with 2035 targets including a 95% reduction in deaths, 90% reduction in TB incidence to 10/100,000 population, and zero families facing catastrophic TB costs (10). Similarly, the Sustainable Development Goals target a 90% reduction in TB deaths and 80% reduction in incidence by 2030 (11). The World Health Organization (WHO) issued The End TB strategy with similar 2035 targets as the Partnership Plan and adds 2025 targets of 90% treatment coverage, 90% treatment success, 90% prevention coverage, and 90% new drugs uptake (10, 12). Achieving the global HIV and TB targets will have a major impact on both epidemics, however, the lack of a comprehensive integrated disease control strategy jeopardizes success for either individual disease effort. In this paper we examine global HIV and TB policy, HIV and TB disease control budgets, and a new combined TB and HIV care continua.

### HIV and TB Policy

In 2004, WHO released guidance on TB-HIV collaborative activities and in 2007 introduced the *Three I’s for TB/HIV* (intensified TB case finding, IPT and infection control) (13–15). Although most high burden countries have adopted the *Three I’s for HIV/TB*, availability of IPT and infection control is limited in most settings (16, 17). HIV is the strongest risk factor for TB, ART and IPT together can reduce TB by nearly 90%, and multiple studies have found that providing early ART for people diagnosed with TB disease can reduce mortality by 50% (4, 7, 8). The 2013 WHO guidelines deferred recommending using earlier ART to prevent TB recommending instead that all HIV-positive TB patients initiate ART (19). The 2015 WHO antiretroviral guidelines finally recommended that ART be initiated irrespective of CD4 cell count with many countries reporting their intention or move to “test and treat” (18). A review of published national HIV policies shows that as of June 2017, 48 countries have published guidelines recommending offering immediate treatment irrespective of CD4 cell count (63% of 2015 HIV burden) (16).

The new science and improved policies has supported remarkable progress over the last decade with an estimated 43 million lives saved, 47% reduction in TB mortality due to TB treatment, and a 32% reduction in HIV-associated TB deaths with the additional benefit of preventing millions of HIV and TB illnesses, deaths and new infections between 1996 and 2014 (1, 6, 20, 21). However, implementation of many TB and HIV policy recommendations is incomplete (16, 17, 22)—in 2015 TB killed one out of three PLHIV globally and it remains the leading cause of morbidity and mortality among PLHIV (1).

### Global investment in HIV and TB control

The goal of controlling the HIV epidemic is now within reach. The President Emergency Plan for AIDS Relief (PEPFAR) and the Global Fund (GF) have focused efforts on jointly addressing HIV-associated TB through HIV and TB single concept notes and promoting joint TB and HIV programming (Table 1) (23, 24). Beginning in 2003, available international and domestic resources for TB and HIV-associated TB averaged of $2.6 billion per year with a cumulative total of 39 billion dollars (HIV control resources not included). For 2012-2016, the budgeted funds have not significantly increased and the average annual resources available was 4 billion. The resource needs estimates for the 2016-2020 Global TB Plan are 13 billion United States Dollars (USD) per year or 65 billion for the five-year period (56 billion for programs and 9 billion for research and development). Since 2003, PEPFAR has invested 1.5 billion USD to address HIV-associated TB (not including resources for HIV control efforts or the PEPFAR contribution to the GF) (23, 25). The GF has invested 5.5 billion USD in TB and an estimated 835 million in HIV/TB related programs (26). Tracking reported international and national resource allocation is important, however, for PLHIV the actual disbursement of funds to support service delivery is critical and, in some settings, does not match reported available budgets. Between 2005 and 2015, these investments have contributed to saving an estimated 9.6 million lives among PLHIV(1). However, despite the 39 billion dollar investment in TB and HIV-associated TB, there are still 400,000 HIV-associated TB deaths annually and TB incidence remains level in most setting (1).

**Table 1:**
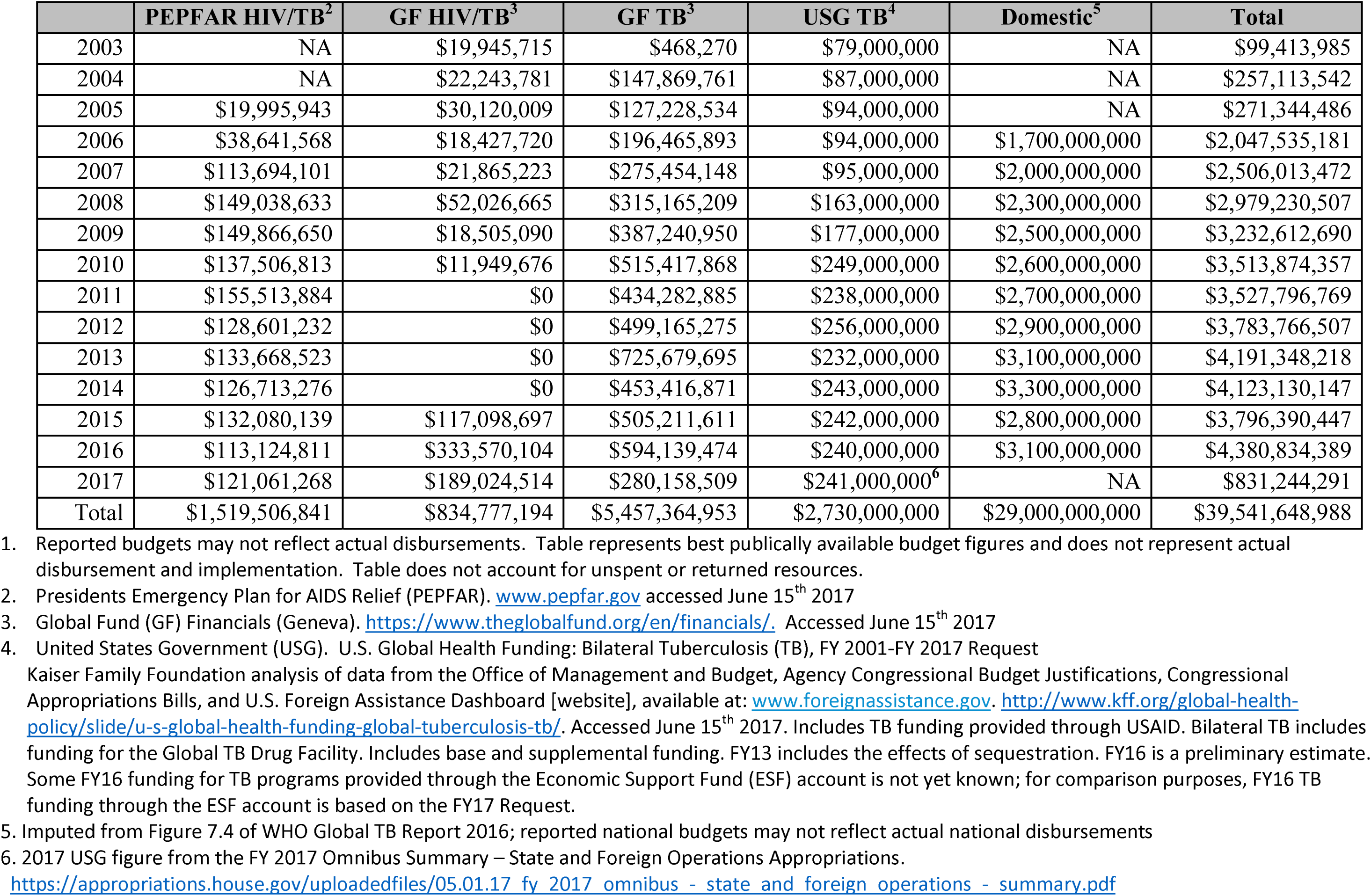
Global TB and HIV Reported Budgets for TB and HIV-associated TB: PEPFAR, GF, USG and Domestic (2003-2017) ^1^.

### HIV, TB and the 90-90-90 Target

Achieving the HIV 90-90-90 target of 73% viral suppression is unlikely without addressing TB and HIV-associated TB. Earlier HIV diagnosis, as part of reaching 90-90-90, allows for earlier ART and IPT to reduce HIV and TB-related morbidity, mortality, and transmission. Early HIV diagnosis also improves access to other interventions including TB screening and measures to prevent nosocomial TB transmission. Although hundreds of millions of HIV tests have been conducted, the yields in many settings have often been below 5% (25). In sub-Saharan Africa, where the global HIV burden is heaviest, HIV prevalence among TB patients ranges from 20-70% and HIV prevalence among persons with presumptive TB (i.e. patients with TB symptoms but who may have a different diagnosis) is 40-80% (27–29). Achieving 90-90-90 will require access to routine HIV testing for individuals at TB clinics, inpatient or outpatient departments and as part of community health service delivery.

Conversely, TB case detection also plays a critical role as a platform for HIV testing and treatment to achieve 90-90-90 target viral suppression. TB incidence is decreasing in places with improve ART coverage, however, TB remains an important means of access ART in countries that do not recommend “test and treat”. Despite its critical importance, TB case detection is only 48% in sub-Saharan Africa where 75% of PLHIV live and ranges from as low as 15% for Nigeria to 96% for Sao Tome and Principe (1); case detection among PLHIV is frequently lower as they often have extra-pulmonary or smear negative disease. MDR-TB case detection is 30% and there is no nationally reported data on the proportion of MDR-TB cases diagnosed with HIV (1). TB screening does not guarantee completion of TB diagnostic evaluation pathway and several studies suggest that only 15–25% of presumptive TB patients receive a complete diagnostic evaluation (30). Expanding access to both HIV and TB screening including the use of rapid diagnostic tests like Xpert MTB/RIF as provides an excellent opportunity to improve identification of people who have HIV and/or TB. Although Xpert MTB/RIF was recommended in 2010 (31) and UNITAID, PEPFAR and GF allocated millions of dollars for implementation, there is no data regarding the proportion of PLHIV who have access to Xpert supported diagnosis. Undiagnosed TB cases are also often undiagnosed PLHIV and represent a significant missed public health opportunity to start people on both HIV and TB treatment as part of achieving 90-90-90 and the TB targets.

As part of achieving the 90-90-90 target, diagnosing HIV-associated TB allows for immediate ART and TB treatment. However, being diagnosed with HIV-associated TB does not guarantee access to life-saving HIV treatment. Globally, in 2015, only 488,364 (79%) of notified HIV positive TB patients were on ART (1), representing a serious service delivery failure for patients who are already in the health care system. This represents only 42% of the estimated 1.2 million PLHIV who developed TB in 2015 (1). Mortality during the first year after TB diagnosis can be as high as 26% (32), however, early diagnosis and combined HIV and TB treatment can reduce mortality by 50% (7, 8). Global TB treatment success in 2014 was 83%, however, it is 75% for PLHIV (1). Reasons for the poorer outcomes include late access to HIV testing and treatment since most countries, until recently, were following WHO recommendations to wait for severe immune degradation before starting HIV treatment (16). An integrated approach to improving earlier HIV testing and TB case detection can synergize and amplify efforts to reach 90-90-90 and TB control targets.

Undiagnosed and untreated TB causes viral replication directly compromising the 90-90-90 durable viral suppression target (33). Routine TB control activities including case finding, provision of treatment or IPT as needed, and TB infection control all contribute to achieving viral load suppression. Integration of TB diagnosis and HIV viral load measurement using a common laboratory platform has significant promise if its costs can be reduced. Achieving both the 90-90-90 HIV target and global TB strategy goals (9–12) are critical to prevent millions of preventable TB and HIV illnesses, deaths and new infections.

### Monitoring and evaluating for “one patient two diseases”

Despite the fact that in many high burden settings up to 71% of people with active TB are co-infected with HIV and up to 89% of PLHIV are infected with *M. tuberculosis* (1, 34), many national health programs rely on entirely separate HIV and TB disease control programs. While allowing single-disease focus, this requires costly duplication of efforts for under-resourced weak health systems. This artificial separation requires patients to shuttle back and forth between programs to the detriment of the health and financial well-being of the individual, community and health system.

Each program has separate monitoring and evaluation efforts focused on monitoring only the HIV or the TB elements of service delivery. Although there may be some overlap such as HIV testing among diagnosed TB patients or TB screening among PLHIV, the monitoring and evaluation system usually neglects the larger monitoring and evaluation perspective needed to control and eliminate both HIV and TB. The common HIV-associated TB care continua reflect a narrow curative approach and focus on what happens once someone is diagnosed with TB. The curative emphasis neglects the prevention impact of interventions that reduce the risk of TB for people infected with HIV and/or *M. tuberculosis*. Similarly, HIV care continua have moved to include accountability from the time of HIV diagnosis to viral suppression. Comprehensive monitoring of care continuum starting “upstream” with HIV and/or *M. tuberculosis* infection has the potential to re-focus efforts and accountability on achieving larger epidemic control progress through both prevention and treatment efforts. Under a more comprehensive monitoring and evaluation framework focused on prevention of TB and/or HIV disease, each person with diagnosed with TB who learns of their HIV status or someone living with HIV who is not on ART and develops TB could be considered a program and public health failure.

### “What is not measured is not done”: combined HIV and TB control care continua

Starting immediate ART irrespective of CD4 cell count can reduce HIV-related morbidity, mortality, transmission and costs (5, 20, 35, 36). However, to realize the benefits PLHIV must optimally engage at each step along the ‘HIV care continua’ (referred to as *continua* hereafter) (37, 38), from early HIV diagnosis to access to ART and viral suppression. HIV care continua have become vital tools to monitor the HIV programmatic efforts and guide actions to achieve 90-90-90-90 (9, 38). As of 2015, only 60% of PLHIV globally had been diagnosed, 46% were reported on ART and 38% PLHIV were estimated to be virally suppressed (39). From the TB care continuum perspective, global TB case detection was only 59%, treatment success was 83%, and only 42% of the estimated people living with HIV and TB were reported to be on dual treatment (1). Achieving HIV 90-90-90 and 90% TB case detection and treatment success by 2020 will require a new combined continuum of care for both preventing and treating HIV and TB (Figure 1). Specifically, the combined TB and HIV continua will need to start with people estimated to be infected with HIV and with *M. tuberculosis*. Instead of focusing on downstream clinical interventions, the comprehensive care continuum would set targets and monitor actions designed to prevent the development of HIV and/or TB disease, provide life-saving treatment, and reduce transmission for both diseases. A comprehensive integrated TB and HIV cascade would have the added benefit of holding programs accountable for both HIV and *M. tuberculosis* infection case detection, prevention of HIV and TB illness, and viral suppression and TB treatment success as surrogates for dual disease control objectives.

**Figure 1:**
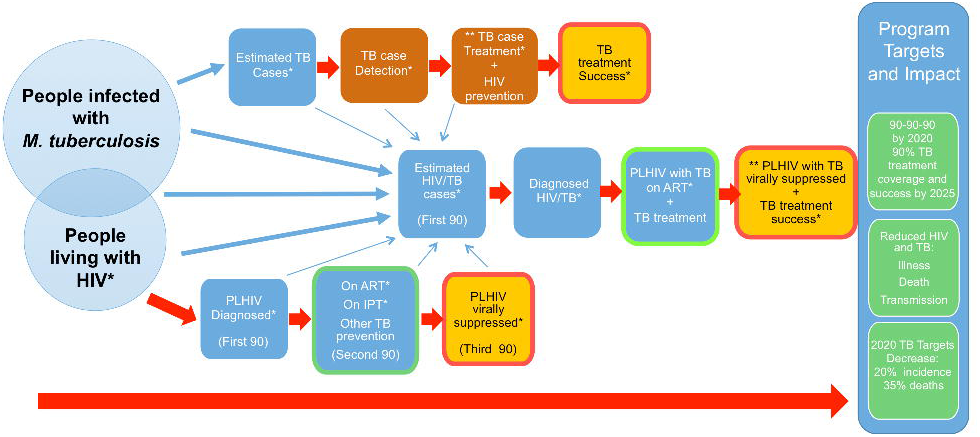
Combined HIV and TB cascade. **Notes:** * Table indicators ** Potential new composite indicator Red outline: individual and public health goal that focuses on dual HIV and TB targets Green outline: programme objective that measures access to HIV and TB interventions

### Comprehensive HIV and TB continua data

Data for a combined HIV and TB continua are available in the public domain, however, they are often fragmented, from different sources, focused on HIV or TB only, and not aggregated into a comprehensive HIV and TB continua. We used data from the *HIV90-90-90watch* website (38, 40), UNAIDS Aidsinfo, and the WHO 2016 Global TB Report (1) for the 22 WHO high TB and TB/HIV burden priority countries (81% HIV/TB burden) to determine the feasibility of constructing comprehensive HIV and TB continua (Table 2). The combined continuum (constructed for South Africa and India) includes new combined integrated indicators that capture PLHIV receiving ART and TB prevention interventions and those with TB who have received successful TB treatment and viral suppression (Figures 2 and 3). These composite indicators further emphasize an integrated approach to HIV and TB control. Viral load data was not available for 13 countries and none of the countries included information regarding the combined indicators: TB treatment success and viral suppression among HIV/TB or TB prevention and ART for PLHIV. PLHIV on ART ranged from 9-70% and TB treatment coverage was between 15-87% (Figure 4). Viral suppression of PLHIV ranged from 38-63% [79-90% among PLHIV on ART] while TB treatment success ranged from 34-94% (Figure 5, Table 2). Only 3% of PLHIV in the 22 countries reported course of IPT.

**Table 2:** HIV, HIV/TB and TB care continua in 22 priority countries for TB and HIV/TB.

| COUNTRY | ART Eligibility | Estimated PLHIV | PLHIV diagnosed | PLHIV on ART | On IPT | PLHIV with viral suppression |  | Estimated annual HIV/TB cases | HIV/TB patients diagnosed | HIV/TB patients on ART | TB treatment success rate among HIV+ TB cases | Estimated TB cases | New and relapse cases notified (TB treatment coverage) | TB treatment success among notified cases |
| --- | --- | --- | --- | --- | --- | --- | --- | --- | --- | --- | --- | --- | --- | --- |
|  |  |  | Number (%) | Number (%) |  | Number (%) | % of those on ART | Number | Number | (%, 2014) |  |  |  | (%, 2014) |
| Angola | <500 | 315,394 |  | 90,204 (29%) |  |  |  | 28,000 | 1,558 (6%) |  |  | 93,000 | 59,705 (64%) | 34% |
| Brazil | All | 827,000 | 715,000 (86%) | 455,000 (55%) |  | 410,000 (50%) | 90% | 13,000 | 9,069 (70%) | 2,852 (22%) | 49% | 84,000 | 73,221 (87%) | 71% |
| Central African Republic | <500* | 118,776 |  | 28,303 (24%) |  |  |  | 8,600 | 1,963 (23%) |  | 68% | 19,000 | 10,459 (55%) | 70% |
| China | <500* | 850,000 | 577,423 (68%) | 386,756 (46%) |  | 328,743 (39%) | 85% | 15,000 | 10,034 (67%) | 3,750 (25%) | 86% | 918,000 | 798,439 (87%) | 94% |
| Congo | <350* |  |  |  |  |  |  | 6,400 | 479 (7%) | 164 (3%) |  | 18,000 | 9,937 (57%) | 69% |
| Democratic Republic of Congo | <500 | 374,097 |  | 122,268 (33%) |  |  |  | 39,000 | 7,154 (18%) | 4,776 (12%) |  | 250,000 | 119,213 (48%) | 89% |
| Ethiopia | <500 | 729,515 |  | 373,933 (51%) | 17,585 |  |  | 16,000 | 8,625 (54%) | 6,848 (43%) |  | 191,000 | 135,951 (71%) | 89% |
| India | <500 | 2,116,581 | 1,523,481 (72%) | 919,141 (43%) |  | 818,035 (39%) | 89% | 113,000 | 44,652 (40%) | 40,925 (36%) | 76% | 2,840,000 | 1,667,136 (59%) | 74% |
| Indonesia | <350 | 691,076 | 177,754 (26%) | 63,066 (9%) | 591 |  |  | 78,000 | 3,523 (5%) | 757 (1%) | 56% | 1,020,000 | 328,895 (32%) | 84% |
| Kenya | <500 | 1,366,771 |  | 857,472 (63%) | 85,392 | 726,370 (53%) | 85% | 36,000 | 26,288 (73%) | 25,030 (70%) | 82% | 107,000 | 81,292 (76%) | 87% |
| Lesotho | <500 | 315,002 |  | 146,790 (47%) |  |  |  | 12,000 | 5,258 (44%) | 4,152 (35%) | 69% | 17,000 | 7,594 (45%) | 70% |
| Liberia | <500 | 30,205 |  | 7391 (24%) |  |  |  | 1,800 | 548 (30%) | 154 (9%) |  | 14,000 | 5,814 (42%) | 74% |
| Mozambique | <350 | 1,505,910 |  | 802,659 (53%) | 130,420 |  |  | 79,000 | 29,827 (38%) | 27,417 (35%) |  | 154,000 | 58,344 (38%) | 89% |
| Myanmar | <500 | 224,794 | 138,370 (62%) | 106,490 (47%) | 3,361 | 92,449 (41%) | 87% | 17,000 | 7,918 (47%) | 3,034 (18%) | 70% | 197,000 | 138,447 (70%) | 87% |
| Namibia | <500 | 229,631 |  | 161,785 (70%) |  | 143,989 (63%) | 89% | 4,900 | 3,796 (77%) | 3,480 (71%) | 80% | 12,000 | 9,614 (80%) | 87% |
| Nigeria | <500 | 3,500,000 |  | 828,867 (24%) | 40,855 |  |  | 100,000 | 14,846 (15%) | 11,141 (11%) | 79% | 586,000 | 87,211 (15%) | 87% |
| Papua New Guinea | <350 | 40,148 |  | 21,198 (53%) |  |  |  | 4,900 | 758 (15%) | 494 (10%) |  | 33,000 | 26,347 (80%) | 70% |
|  |  |  | Number (%) | Number (%) |  | Number (%) | % of those on ART | Number | Number | (%, 2014) |  |  |  | (%, 2014) |
| South Africa | <500 | 6,669,360 |  | 3,217,097 (48%) | 409,496 | 2,541,507 (38%) | 79% | 258,000 | 157,505 (61%) | 133,116 (52%) | 76% | 454,000 | 287,224 (63%) | 78% |
| Thailand | All | 437,700 | 389,027 (87%) | 272,750 (62%) |  | 223,372 (51%) | 82% | 15,000 | 7,819 (52%) | 5,389 (36%) | 67% | 117,000 | 62,135 (53%) | 80% |
| Tanzania | <500 | 1,385,780 |  | 740,078 (53%) |  |  |  | 57,000 | 20,117 (35%) | 17,063 (30%) | 87% | 164,000 | 60,895 (37%) | 90% |
| Zambia | <500 | 1,211,880 |  | 758,646 (63%) |  |  |  | 38,000 | 20,967 (55%) | 15,897 (42%) |  | 63,000 | 36,741 (58%) | 85% |
| Zimbabwe | <500 | 1,400,000 |  | 879,271 (63%) | 38,489 | 723,640 (52%) | 82% | 26,000 | 18,072 (70%) | 12,924 (50%) | 68% | 38,000 | 26,990 (72%) | 81% |

\*reported ART eligibility criteria
All data are from 2015, unless mentioned.
PLHIV – people living with HIV, TB – tuberculosis, IPT – isoniazid preventive therapy, ART – antiretroviral treatment

**Figure 2:**
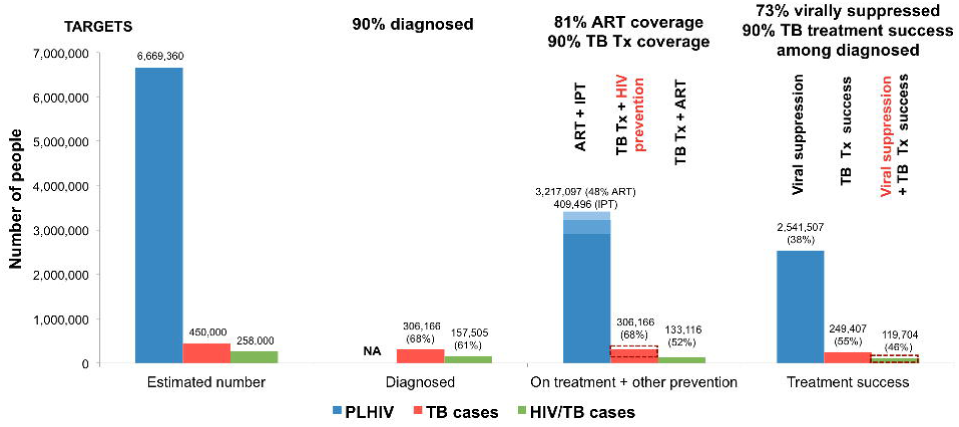
Continua of HIV and TB in South Africa. **Notes:** Percentages are calculated using the estimated number of people as the denominator. TB continuum is from 2014 and HIV and HIV/TB continua are from 2015. HIV targets are for 2020 and TB targets are for 2025. NA – not available, Tx – treatment, ART – antiretroviral treatment, IPT – isoniazid preventive therapy, TB - Data on interventions in red are not collected.

**Figure 3:**
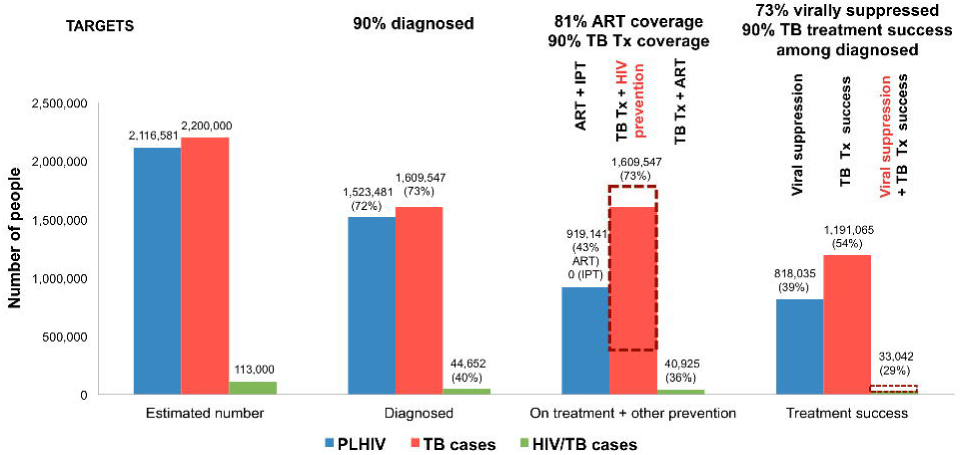
Continua of HIV and TB in India. **Notes:** Percentages are calculated using the estimated number of people as the denominator. TB continuum is from 2014 and HIV and HIV/TB continua are from 2015. HIV targets are for 2020 and TB targets are for 2025. NA – not available, Tx – treatment, ART – antiretroviral treatment, IPT – isoniazid preventive therapy, TB Data on interventions in red are not collected.

**Figure 4:**
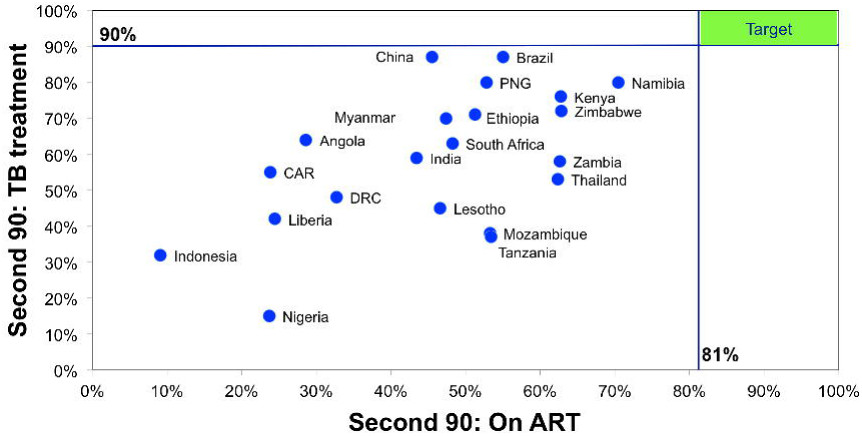
Country progress towards HIV and TB targets: Proportion of PLHIV on ART and estimated TB cases on TB treatment for 22 priority countries. **Notes:** CAR – Central African Republic, DRC – Democratic Republic of Congo, PNG – Papua New Guinea. Data are from 2015 for countries.

**Figure 5:**
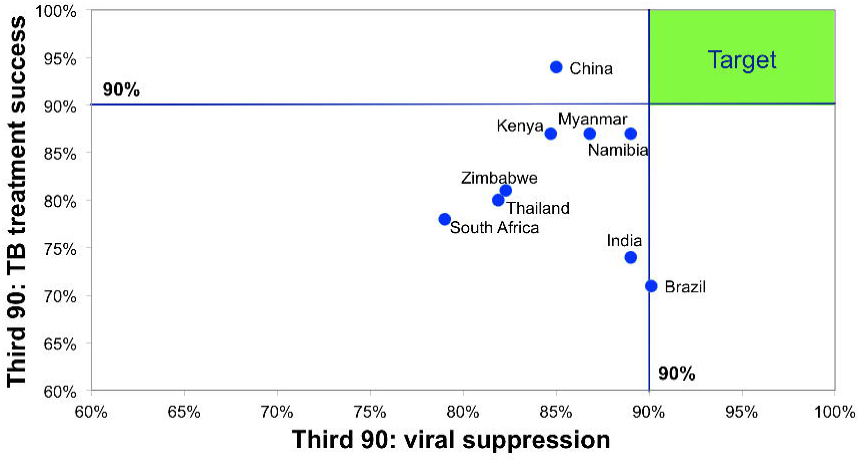
Country progress towards HIV and TB targets: Proportion of PLHIV on ART with viral suppression and TB treatment success among notified cases for 22 priority countries. **Notes:** Data on TB treatment success from 2014 and on viral suppression from 2015

## Discussion

HIV, TB and HIV-associated TB continues to pose a significant public health threat despite a robust global and domestic response including a 39 billion USD global budget for TB and HIV-associated TB for 2003 to 2015. TB and HIV disease control are usually separate endeavors with minimal overlap. The concept of continua of care for HIV, TB and HIV-associated TB is not new, however its potential has only been realized for one or the other disease in a limited number of settings. Specifically, available HIV continua data demonstrate that countries are moving toward 90-90-90 (38, 40). Similarly, national and sub-national TB cohort reports document case detection and treatment success (41, 42). Some efforts have been made to monitor the treatment end of the HIV-associated TB care continua including screening for TB disease, testing for HIV and providing ART for PLHIV who have TB disease (1). There are some TB and HIV continua data for the 22 high burden countries. However, unsurprisingly, no high burden countries reported comprehensive combined 90-90-90 and TB continua data. While most HIV and TB programs report shared indicators such as ART for TB cases, none report integrated comprehensive continuum that include HIV diagnosis and/or *M. tuberculois* screening, ART and/or IPT, TB and HIV associated TB case detection and treatment, viral suppression and TB and treatment outcomes. The TB specific outcomes for HIV-associated TB treatment are also unavailable for most countries. It is also not clear from most of the available data whether people diagnosed with HIV and TB were already on IPT and/or ART. Although seemingly a minor detail, both the proportion of new HIV diagnoses among people living with TB and the proportion of new TB cases among PLHIV not on treatment could also become critical measures to assess the success of HIV and TB prevention efforts.

Although the TB monitoring and evaluation system with its reliance on quarterly cohorts is over 40 years old, the HIV care continuum is a relatively new concept (37) and many countries are still struggling to measure continua and the 90-90-90 target (38). For example, only a few countries could produce national cohorts and/or national program data that include HIV diagnosis through viral suppression (38). In contrast toTB cohort analyses based on following individual patients over time, most countries derive indirect ross-sectional estimates of HIV continua numerators, which may be prone to data errors and methodological flaws. Specifically, without patient identifiers and a cohort system, patient loss after diagnosis, duplication of patients in the system, and lack of individually linked viral load results may compromise the validity of some continua. Also, this study did not evaluate the availability of high burden sub-national or population-specific care continuums (e.g., key populations) for HIV or TB, which may more directly inform strategies for targeted program improvement and community incidence interruption.

## Conclusion

The adoption of the 90-90-90 target has significant promise for ending the HIV epidemic. Focusing on diagnosing and ensuring that PLHIV access earlier ART also makes sense from a TB control perspective since it is also proven to prevent HIV-associated TB disease, deaths, and transmission. Likewise, diagnosing and treating active TB can be life-saving, prevents transmission and, for PLHIV, allows for achieving HIV viral suppression. Given the significant interaction between the two pathogens, combining disease control objectives to achieve HIV 90-90-90, end of AIDS, and the TB 90-90-90 targets makes sense. In geographic settings and populations with high TB and HIV burdens, changing health services so that they can deliver a comprehensive combined TB and HIV continuum of care will be essential to achieving the 90-90-90 and TB control targets. Simplifying and combining the existing multiple, separate disease control strategies, reducing inefficient, duplicative siloed funding and programming, and combining and simplifying monitoring and evaluation of service delivery and impact are essential given current resource constraints. The goal of both HIV and TB programs, to control and eliminate both diseases, can only be accomplished when both TB and HIV targets are achieved. Using a comprehensive HIV and TB care continuum provides an opportunity to harness both the HIV and TB response to improve the health of people infected with HIV and/or *M. tuberculosis infection*, diagnosed with TB disease, their families and the community.

